# Applying Artificial Intelligence to Identify Common Targets for Treatment of Asthma, Eczema, and Food Allergy

**DOI:** 10.1101/2023.06.29.547141

**Authors:** Bonnie Hei Man Liu, Andre Rayner, Andrew R. Mendelsohn, Anastasia Shneyderman, Michelle Chen, Frank W. Pun

## Abstract

Allergic disorders are common diseases marked by the abnormal immune response towards foreign antigens that are not pathogens. Often patients with food allergy also suffer from asthma and eczema. Given the similarities of these diseases and a shortage of effective treatments, developing novel therapeutics against common targets of multiple allergies would offer an efficient and cost-effective treatment for patients. Herein, we employed the artificial intelligence-driven target discovery platform, PandaOmics, to identify common targets for treating asthma, eczema, and food allergy. Thirty-two case-control comparisons were generated from 15, 11, and 6 transcriptomics datasets related to asthma (558 cases, 315 controls), eczema (441 cases, 371 controls), and food allergy (208 cases, 106 controls) respectively, and allocated into three meta-analyses for target identification. Top-100 high-confidence targets and Top-100 novel targets were prioritized by PandaOmics for each allergic disease. Six common high-confidence targets (i.e., *IL4R*, *IL5*, *JAK1*, *JAK2*, *JAK3*, and *NR3C1*) across all three allergic diseases have approved drugs for treating asthma and eczema. Based on the targets’ dysregulated expression profiles and their mechanism of action in allergic diseases, three potential therapeutic targets were proposed. *IL5* was selected as a high-confidence target due to its strong involvement in allergies. *PTAFR* was identified for drug repurposing, while *RNF19B* was selected as a novel target for therapeutic innovation. Analysis of the dysregulated pathways commonly identified across asthma, eczema, and food allergy revealed the well-characterized disease signature and novel biological processes that may underlie the pathophysiology of allergies. Altogether, our study dissects the shared pathophysiology of allergic disorders and reveals the power of artificial intelligence in the exploration of novel therapeutic targets.

## Introduction

Allergies, particularly in children, play a critical role in the overall health and well-being. The prevalence of food allergy worldwide has risen from about 2% in 1950 to over 7% in 2020(1). In the US alone, over 170 foods are reported to elicit an allergic response(2). Allergic reactions can significantly impact a child’s development and quality of life, with reactions ranging from mild hives to anaphylactic reactions and even death(3, 4). Pathologically, food allergy can be Immunoglobulin E (IgE)-mediated, non-IgE-mediated, or mixed, with the underlying mechanisms for non-IgE-mediated reactions remaining poorly defined. Besides food allergy, asthma and eczema are generally considered as allergic diseases, with the three often coexisting among children globally(5, 6). The term ‘atopic march’ denotes the increased occurrence of allergic disorders, such as asthma and food allergy, after the development of atopic dermatitis(7), suggesting the need for developing treatments that address multiple allergies simultaneously. This approach would enhance patient compliance and make drug development more effective.

Traditional drug discovery approaches are tedious and expensive, often taking 5 to 10 years of intense research and costing millions of dollars with a low probability of success. However, with the availability of large and high-dimensional data derived from multi-omics analysis of healthy volunteers and patients’ tissue samples, as well as data on compounds, publications, grants, and clinical trials, artificial intelligence (AI) has emerged as a powerful technology to automate data analysis and accelerate novel therapeutic target identification(8, 9). In particular, machine learning and deep learning can contribute significantly to the prediction of novel therapeutic targets and the repurposing of existing drugs(10, 11, 12, 13). Applying AI-driven approaches not only significantly reduces the time and cost spent on drug development, but may also improve the success rate of clinical trials to remarkably benefit the patients(14).

In the current study, we used PandaOmics, an AI-driven target discovery platform, as a primary tool to analyze publicly available datasets and search for potential therapeutic targets in asthma, eczema, and food allergy. As shown in Figure 1, transcriptomics datasets for asthma, eczema, and food allergy were retrieved from public repositories and uploaded onto PandaOmics to undergo target identification, differential gene expression, and pathway analyses. Eleven common unique targets were selected across the three indications and further evaluated with their potential in treating these allergies. Among them, *IL5* was selected as a high-confidence target due to its strong involvement in allergies. *PTAFR* demonstrated its promise for drug repurposing, while *RNF19B* offered a high potential for novel therapeutic intervention. This piece of work indicated the potential of AI in driving a rapid target discovery process.

**Figure 1.**
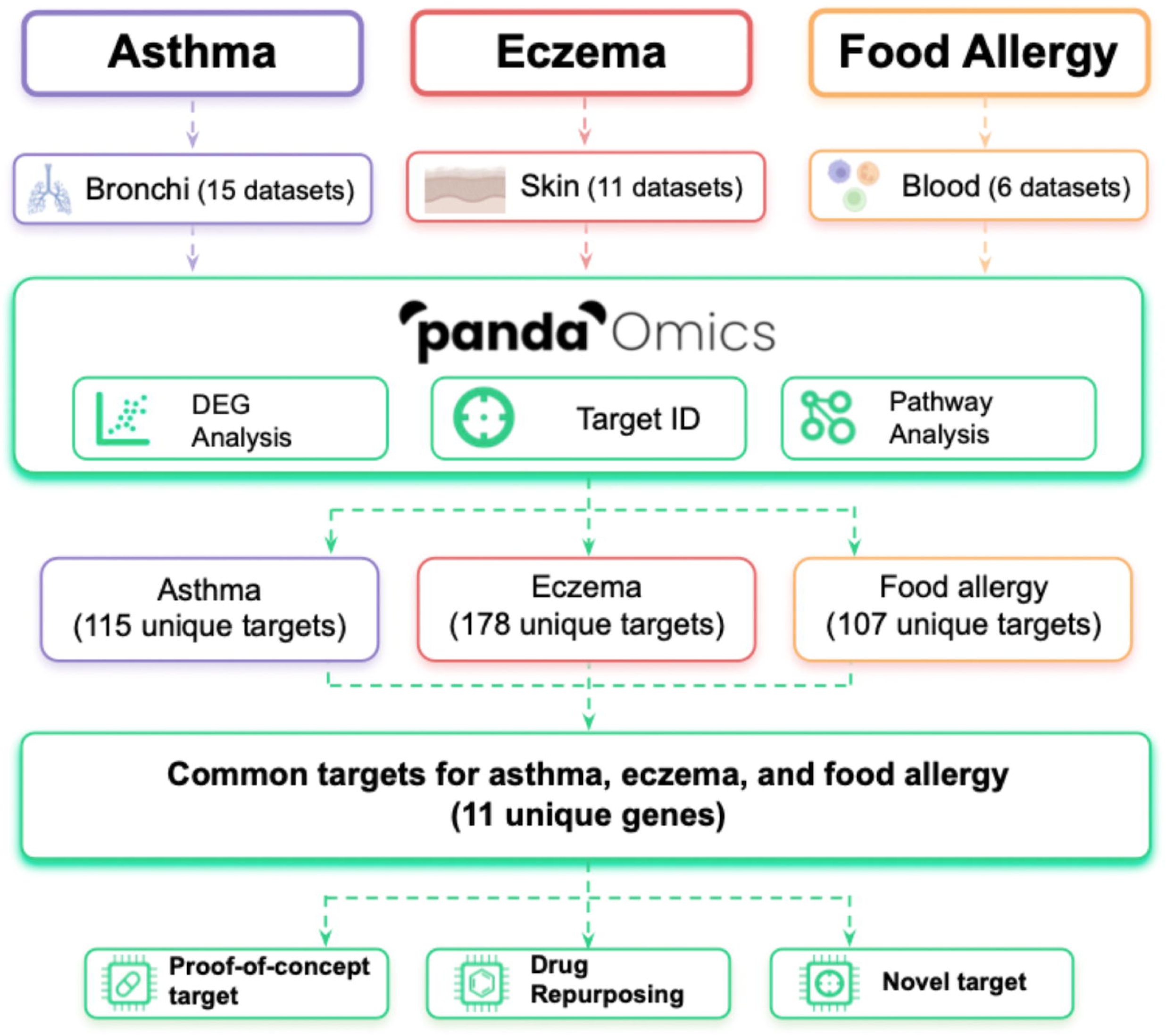
Study scheme of applying PandaOmics to asthma, eczema, and food allergy. Transcriptomics datasets with bronchial epithelial tissue, skin, and blood for asthma, eczema, and food allergy were retrieved from the public repositories. Case-control comparisons were generated from datasets consisting both samples from patients and healthy individuals, and allocated into three meta-analyses based on their disease types in PandaOmics for target identification (Target ID). Expression profiles of the differentially expressed genes (DEGs) and the list of dysregulated signaling pathways were also obtained from PandaOmics for downstream analysis. After target prioritization by PandaOmics under two novelty settings, followed by DEG analysis, 115, 178, and 107 unique dysregulated targets for asthma, eczema, and food allergy were shortlisted. Among the eleven commonly dysregulated targets in the three indications, targets for proof-of-concept, drug repurposing, and novel therapeutic intervention were selected based on consistency of the dysregulated expression across the comparisons in each indication, relevance to allergies, drug availability, and clinical trial status.

## Materials and Methods

### Dataset and comparison selection

Bulk transcriptomics datasets with bronchial epithelial and skin tissues were selected for target identification for asthma and eczema, respectively, while target identification for food allergy was performed using datasets with blood or immune cells. As a result, 15, 11, and 6 relevant datasets were selected for asthma, eczema, and food allergy, respectively. Samples from 5 out of 6 datasets selected for food allergy were collected from children and adolescents, while the remaining one was with a mixture of samples from adolescents and adults. All samples used in the eczema-related datasets were from adults. For asthma, 13 and 1 datasets were generated from samples collected from adults and children, respectively. No age information was retrieved for the remaining dataset for asthma. They were retrieved from Gene Expression Omnibus (GEO) and ArrayExpress databases and processed before being uploaded onto PandaOmics for target identification. One case-control comparison was generated from each of the datasets selected. Case-control comparisons for target identification were listed in Table 1. Normal relevant tissue from healthy individuals was chosen as the controls.

**Table 1.** Transcriptomics case-control comparisons for asthma, eczema, and food allergy.

| Serie | Type | Platform | Tissue Source | Experimental design | # Case | # Control | Year | PMID |
| --- | --- | --- | --- | --- | --- | --- | --- | --- |
| *<u>Asthma meta-analysis</u>* |  |  |  |  |  |  |  |  |
| GSE4302 | Microarray | GPL570 | Bronchial epithelium | Asthma vs Healthy | 42 | 28 | 2006 | 17898169 |
| GSE41649 | Microarray | GPL96 | Bronchial epithelium | Asthma vs Healthy | 4 | 4 | 2012 | 19842841 |
| GSE41861 | Microarray | GPL570 | Bronchial epithelium | Asthma vs Healthy | 51 | 30 | 2012 | - |
| GSE43696 | Microarray | GPL6480 | Bronchial epithelium | Asthma vs Healthy | 88 | 20 | 2013 | 24518246 |
| GSE63142 | Microarray | GPL6480 | Bronchial epithelium | Asthma vs Healthy | 128 | 27 | 2014 | 25338189 |
| GSE64913 | Microarray | GPL570 | Bronchial epithelium | Asthma vs Healthy | 22 | 37 | 2015 | 28045928 |
| GSE89809 | Microarray | GPL13158 | Bronchial epithelium | Asthma vs Healthy | 38 | 18 | 2016 | 28933920 |
| GSE104468 | Microarray | GPL21185 | Bronchial epithelium | Asthma vs Healthy | 12 | 12 | 2017 | 28294656 |
| GSE179156 | Microarray | GPL570 | Bronchial epithelium | Asthma vs Healthy | 38 | 29 | 2021 | 34233672 |
| GSE85567 | RNA-seq | GPL11154 | Bronchial epithelium | Asthma vs Healthy | 57 | 28 | 2016 | 27942592 |
| GSE109484 | RNA-seq | GPL16791 | Bronchial epithelium | Asthma vs Healthy | 33 | 14 | 2018 | 29548336 |
| GSE101720 | RNA-seq | GPL18573 | Bronchial epithelium | Asthma vs Healthy | 14 | 18 | 2017 | 30190271 |
| GSE106388 | RNA-seq | GPL18573 | Bronchial epithelium | Asthma vs Healthy | 15 | 4 | 2017 | - |
| GSE118761 | RNA-seq | GPL11154 | Bronchial epithelium | Asthma vs Healthy | 21 | 29 | 2018 | 32113981 |
| GSE158752 | RNA-seq | GPL18573 | Bronchial epithelium | Asthma vs Healthy | 25 | 17 | 2020 | 32531372 |
| *<u>Eczema meta-analysis</u>* |  |  |  |  |  |  |  |  |
| GSE58558 | Microarray | GPL570 | Skin | Eczema vs Healthy | 18 | 17 | 2014 | 24786238 |
| GSE59294 | Microarray | GPL570 | Skin | Eczema vs Healthy | 16 | 7 | 2014 | 25482871 |
| GSE99802 | Microarray | GPL570 | Skin | Eczema vs Healthy | 59 | 53 | 2018 | 30121291 |
| GSE130588 | Microarray | GPL570 | Skin | Eczema vs Healthy | 51 | 42 | 2019 | 30194992 |
| GSE133385 | Microarray | GPL570 | Skin | Eczema vs Healthy | 30 | 30 | 2019 | 31356921 |
| GSE133477 | Microarray | GPL570 | Skin | Eczema vs Healthy | 79 | 40 | 2019 | 31419544 |
| GSE140684 | Microarray | GPL570 | Skin | Eczema vs Healthy | 31 | 21 | 2019 | 27304428 |
| GSE121212 | RNA-seq | GPL16791 | Skin | Eczema vs Healthy | 21 | 27 | 2019 | 30641038 |
| GSE137430 | RNA-seq | GPL20301 | Skin | Eczema vs Healthy | 40 | 39 | 2020 | 32428528 |
| GSE141571 | RNA-seq | GPL20301 | Skin | Eczema vs Healthy | 39 | 41 | 2020 | 32428543 |
| GSE157194 | RNA-seq | GPL21290 | Skin | Eczema vs Healthy | 57 | 54 | 2020 | 32615169 |
| *<u>Food allergy meta-analysis</u>* |  |  |  |  |  |  |  |  |
| GSE88796 | Microarray | GPL10558 | PBMC* | Food allergy vs Healthy | 65 | 28 | 2016 | 27788149 |
| GSE171660 | Microarray | GPL13607 | Whole blood | Food allergy vs Healthy | 11 | 9 | 2021 | - |
| GSE114065 | RNA-seq | GPL20301 | T cells | Food allergy vs Healthy | 65 | 36 | 2018 | 30120223 |
| GSE165316 | RNA-seq | GPL24676 | B cells | Food allergy vs Healthy | 18 | 9 | 2021 | 34466226 |
| GSE189149 | RNA-seq | GPL24676 | T cells | Food allergy vs Healthy | 30 | 18 | 2021 | - |
| GSE196495 | RNA-seq | GPL16791 | T cells | Food allergy vs Healthy | 19 | 6 | 2022 | 35266148 |

\* PBMC stands for Peripheral Blood Mononuclear Cells.

In addition, GSE41861 and GSE158752 that have information on disease severity, were selected for identifying indicators for disease severity. There were 10, 81 and 30 samples for severe asthma, mild/moderate asthma, and healthy control in GSE41861, while GSE158752 contained 25, 25, and 17 samples isolated from patients with severe asthma, mild/moderate asthma, and healthy individuals. Three comparisons were generated for each dataset, comparing the expression differences between severe asthma and controls, mild/moderate asthma and controls, as well as asthma samples with different severities (S4 Table).

### Target identification by PandaOmics

To perform target identification for the three indications with PandaOmics, the 15, 11, and 6 case-control comparisons for asthma, eczema, and food allergy were allocated into 3 independent meta-analyses, which were shown in Table 1. PandaOmics ranked the targets for each disease with a combination of multiple AI scores derived from omics and text data. These AI and bioinformatic models were validated with the Time Machine approach to ensure the identification of truly novel targets for the disease of interest(15). The Omics AI scores include thirteen models representing the target-disease association based on molecular connections between the proposed target and the selected disease. Most of these scores are calculated in real time according to the input of datasets. On the other hand, Text-based AI scores, KOL scores, and Financial scores are calculated from literature and publicly available databases. Scores of each of the models are given on a normalized scale from 0 to 1, with higher scores corresponding to better target-disease association as predicted by the model. Detailed definitions of the scores were described in the User manual section of PandaOmics (https://insilico.com/pandaomics/help). The Metascore (final ranking) provided a list of targets/genes along with the disease of interest for human evaluation and consideration.

To generate therapeutic targets with various degrees of novelty, two settings of filters and scores were employed to obtain a list of targets with high confidence and novelty, respectively. Targets with high confidence in the allergy indications were prioritized based on the Omics, Text-based, Financial, and KOL scores. On the other hand, only the Omics scores were employed for novel target identification. Targets belonging to druggable classes and not regarded as essential genes according to the Therapeutic Target Database (TTD) were selected for further evaluation to avoid potential toxicity.

### Pathway analysis

The assessment of signaling pathway activation status was performed by the proprietary single network model iPANDA incorporated in PandaOmics(16). In general, iPANDA utilized the data from differential gene expression and pathway topology decomposition to rapidly identify sets of biological relevant pathway signatures with notable noise reduction. The Reactome database serves as the basis of the hierarchical organization of signaling pathways(17). Pathway activation or suppression was indicated by the positive or negative iPANDA score, respectively.

### Statistical analysis

Differential expression analysis was performed using the eBayes function in Limma package(18). Gene expression changes among the healthy cohort and different severity groups of asthma patients were plotted on box-plots, and two-tailed t-test was used to estimate the statistical significance.

## Results

### Biological processes are commonly dysregulated in allergies

In view of the interconnection between asthma, eczema, and food allergies, dysregulated signaling pathways in the case-control comparisons for the three diseases were compared to determine the commonly dysregulated mechanisms for allergies using the iPANDA algorithm. Pathways were organized based on the hierarchical structure provided by the Reactome database, with each of them corresponding to one top-level pathway. One hundred and thirty-one signaling pathways were identified as unidirectionally dysregulated in all three diseases, with 104 being activated and 27 being inhibited. The biological processes with the highest number of activated pathways included signal transduction (17%), gene expression (transcription) (13%), immune system (13%), and metabolism of proteins (10%) (Fig. 2A). The details of dysregulated pathways were listed in S1 Table.

**Figure 2.**
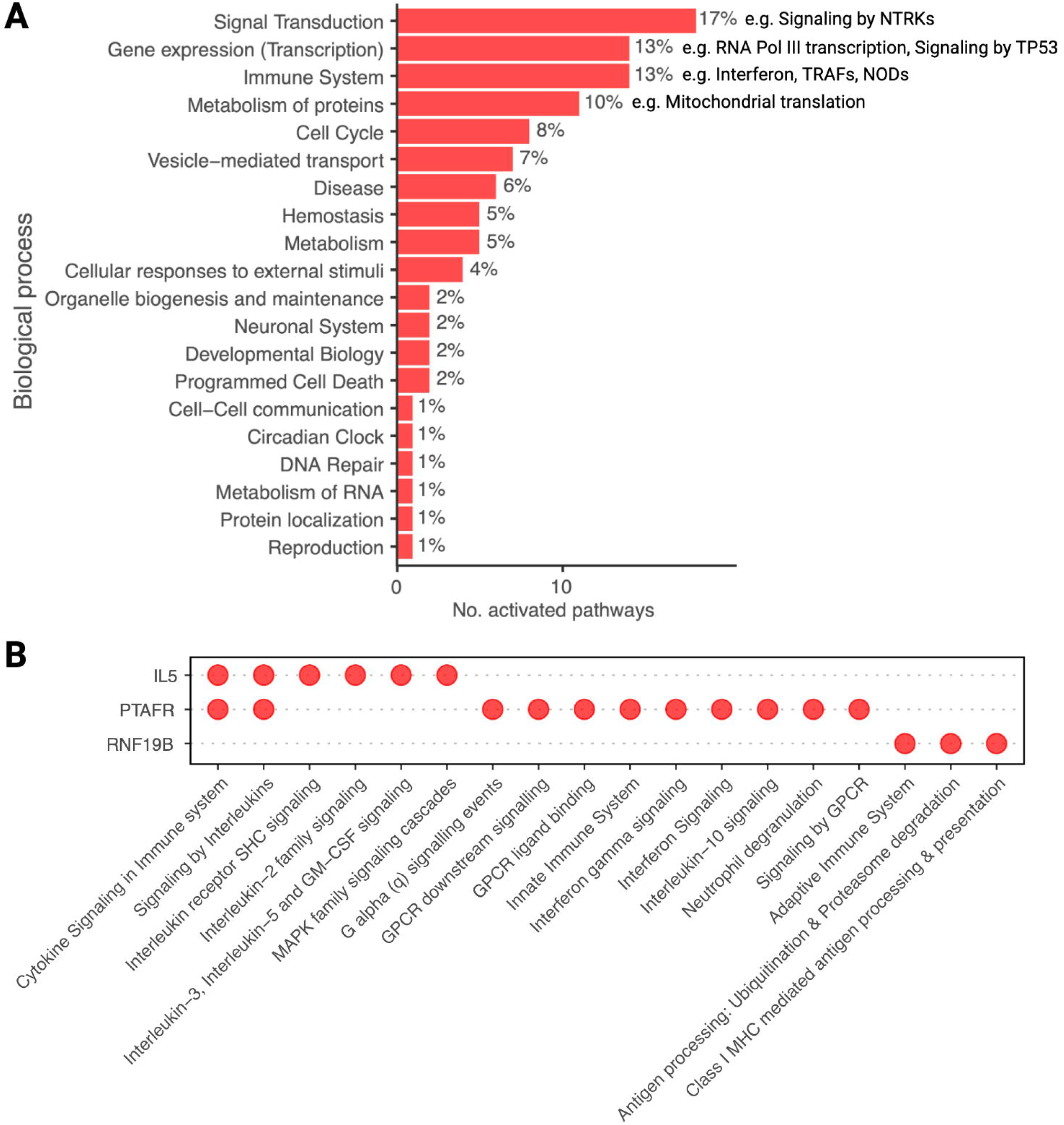
Common activated pathways across asthma, eczema, and food allergy. (A) One hundred and four common activated pathways in asthma, eczema, and food allergy were categorized into twenty biological processes. The percentage of activated pathways in a particular process over all activated pathways was shown next to each bar. Please note that one pathway belonged to two independent biological processes. (B) Common activated pathways associated with IL5, PTAFR, and RNF19B in asthma, eczema, and food allergy were displayed in the dot plot. Presence of a red dot indicated the association between the given target and the pathway.

### Targets associated with FDA-approved drugs for three allergy indications are highly ranked in PandaOmics

To understand the effectiveness of PandaOmics in revealing potential therapeutic targets for allergies, the United States Food and Drug Administration (FDA)-approved drugs for asthma, eczema, and food allergy, and their associated targets were retrieved via literature search and clinical trial result evaluation. No drug has been approved for food allergy by the FDA. Eighteen targets were associated with twenty-six compounds approved for treating asthma and eczema (Table 2). The ranking of these targets under the high confidence setting for each indication in PandaOmics was stated in Table 2, with six of them (i.e., *IL4R*, *IL5*, *JAK1*, *JAK2*, *JAK3*, and *NR3C1*) ranked as the Top-100 targets across all three allergic diseases. *IL13* and *TYK2* were also top-ranked in eczema and food allergy, and asthma and eczema, respectively.

**Table 2.**
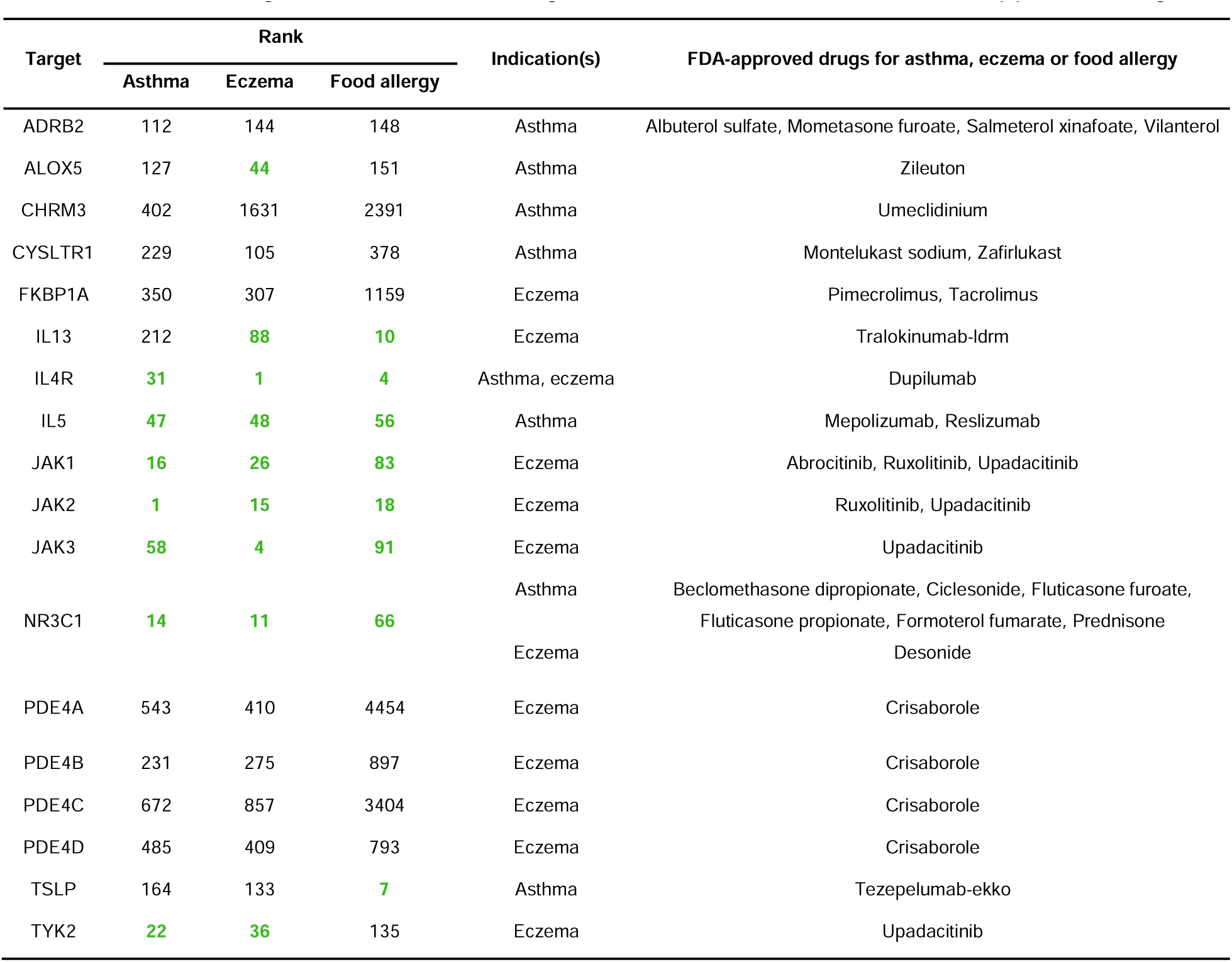
A list of targets across three allergic diseases with associated FDA-approved drugs.

Even though most of these drugs were approved for only one specific indication, given that the target is often found in other allergic indications, one could further evaluate the repurposing of these drugs. Besides, the successful retrieval of the known therapeutic targets for allergies indicated the feasibility of identifying targets with adequate therapeutic value using PandaOmics.

### Interleukin-5 - a high-confidence therapeutic target for allergies

Top-100 high-confidence targets and Top-100 novel targets were prioritized by PandaOmics for each allergic disease. Expression profiles of these targets in allergic patients and healthy individuals were further analyzed, giving 115, 178, and 107 unique dysregulated targets for asthma, eczema, and food allergies, respectively (Fig. 1). Eleven targets (i.e., *CLOCK*, *DCTPP1*, *IFNG*, *IL5*, *PSMA2*, *PSMA4*, *PTAFR*, *RNF19B*, *TGFB1*, *UBE2F*, and *VDR*) were commonly dysregulated across three tested allergy indications (Fig. 3A). All of them were upregulated in the allergies when compared with the healthy controls. Amongst these targets, Interleukin-5 (*IL5*) was selected as a promising high-confidence therapeutic target for allergies. It is a pathogenic target frequently reported in the tested allergy indications(19, 20, 21, 22, 23, 24). Its upregulation was detected in 13 out of 15, 10 out of 11, and 5 out of 5 comparisons for asthma, eczema, and food allergy (S2 Table). *IL5* ranked 54^th^, 68^th^, and 56^th^ in the meta-analysis of asthma, eczema, and food allergy, respectively. Out of the thirteen Omics models, five (i.e., Knockouts, Pathways, Relevance, Heterogenous graph walk, and Matrix factorization) scored 0.8 or above, meaning *IL5* was prioritized by the corresponding models across the tested indications (Fig. 3B).

**Figure 3.**
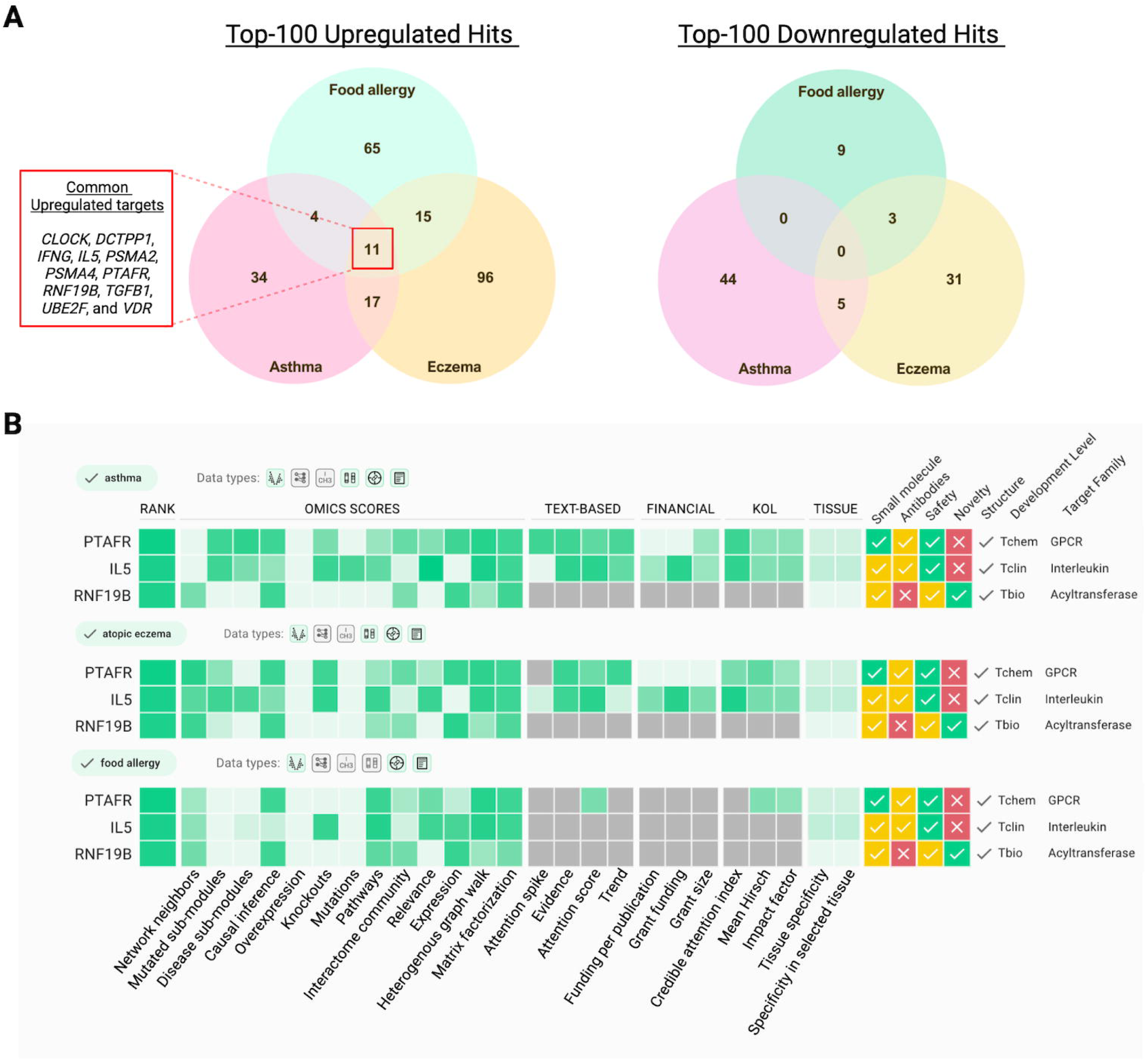
PandaOmics-derived targets matched up against each other for comparison. (A) Two lists of Top-100 targets with high confidence and novelty for each indication were obtained from PandaOmics. Based on the consistency of the dysregulated expression across the comparisons in each indication, as well as the statistical significance, these 200 unique targets were classified into three groups: upregulated (*left*), downregulated (*right*), and no difference (not shown). Upregulated targets in the three indications were overlapped to identify the shared targets. The same applied to the downregulated targets. Eleven targets were commonly activated across the three allergic indications. (B) Screenshot of the Target ID page of PandaOmics for the meta-analyses for asthma, eczema, and food allergy. *IL5*, *PTAFR*, and *RNF19B* were revealed as the target for proof-of-concept, drug repurposing, and novel therapeutic design, respectively.

Through mediating the maturation, proliferation, activation, and migration of eosinophils via JAK/STAT, NF-κB, and PI3K pathways(25), IL5 serves as an influential pro-inflammatory cytokine. In asthma, IL5 coordinates type 2 inflammation, a phenotype observed in 50-70% of asthma patients(26). Its dysregulation is also implicated in airway remodeling, such as increasing airway fibrotic response(27) and thickening of the reticular basement membrane(19). It is noteworthy that three antibody inhibitors targeting IL5/IL5R signaling, i.e., mepolizumab, reslizumab, and benralizumab, have been approved for the treatment of asthma. They all offered excellent safety and efficacy profiles in the clinical trials for asthma. The efficacy of these inhibitors in treating food allergy is yet to be determined.

### *PTAFR* - a high-confidence target for drug repurposing in allergies

Beside *IL5*, we selected Platelet Activating Factor Receptor (*PTAFR*) as a potential therapeutic target for drug repurposing (Fig. 3B). PTAFR was ranked as the 1st, 60th, and 35th target in the meta-analysis of asthma, eczema, and food allergy. It performed well (scored ≥ 0.8) in Causal Inference, Heterogenous graph walk, and Matrix factorization models in the three meta-analyses (Fig. 3B). It was significantly overexpressed in patient samples in 5, 10, and 1 comparisons for asthma, eczema, and food allergy, respectively (S2 Table). Functionally, the binding of platelet-activating factor to PTAFR activates PKC and calcium release, which subsequently induces platelet aggregation, leukocyte reactivity, vasodilation, and inflammation(28). Multiple lines of evidence suggest the potential of inhibiting PTAFR and its signaling to suppress allergic response and inflammation(29, 30, 31). For instance, PTAFR-deficient mice displayed a reduced level of anti-ovalbumin IgE associated with attenuated allergic markers when compared with wild-type mice upon challenges with ovalbumin diet(32). Furthermore, rupatadine, a small molecule inhibitor of PTAFR and histamine H1 receptor(33), is indicated for allergic rhinitis with good long-term safety and tolerability(34, 35). Given it has not been tested in the three indications and its target *PTAFR* was a commonly upregulated gene, rupatadine could be a prospective repurposing candidate for asthma, eczema, and food allergy.

### *RNF19B* - an unreported target for novel therapeutic intervention in allergies

*RNF19B*, a E3 ubiquitin-protein ligase, was ranked as the Top-20 target and scored ≥ 0.8 in the two Omics models (i.e., Causal Inference and Expression) for the three indications (Fig. 3B). In particular, its overexpression reached statistical significance in all comparisons for eczema (*p* < 0.05) (S2 Table). RNF19B ubiquitinates URKL-1 protein for degradation(36). Despite the fact that there are limited amounts of functional studies for RNF19B, the existing evidence highlights its immunological role by reporting its pivotal contribution to natural killer cell cytotoxicity(37, 38). It is suggested to be a positive regulator of STAT1 and NFκB signalings to mediate innate immunity(39, 40). RNF19B-knockout mice exhibited reduced inflammation, less cytokine production, and defective neutrophils and macrophages during *Streptococcus pneumoniae* infection(41). Considering that allergic inflammation is a hallmark of the three analyzed indications(42) and the strong association between RNF19B and immune regulation, these findings indicate the likelihood of RNF19B involvement in allergic diseases.

### Common activated pathways in allergies associated with *IL5*, *PTAFR*, and *RNF19B*

With the aid of the iPANDA algorithm, dysregulated pathways associated with *IL5*, *PTAFR,* and *RNF19B* were also evaluated to dissect their potential roles in allergy (Fig. 2B, S3 Table). The majority of the dysregulated pathways linked to these targets belonged to the immune system. In general, the dysregulated pathways associated with *IL5*, *PTAFR*, and *RNF19B* did not overlap, indicating these targets should have different roles in allergies. Our results were in accordance with the literature and supported the potency of *IL5, PTAFR,* and *RNF19B* as therapeutic targets for allergies.

### Upregulation of *PTAFR* could serve as an indicator of disease severity in asthma

Besides exploring the expression profiles in the primary affected tissue, we examined the association between disease severity and the dysregulation of *IL5*, *PTAFR,* and *RNF19B*. Information on disease severity was available in some of the asthma-related datasets; therefore, GSE41861 and GSE158752 were utilized for such analysis. Apart from being significantly overexpressed in asthmatic bronchial epithelial tissues compared to the controls, *PTAFR* was particularly upregulated in samples from severe asthma compared to those with mild/moderate asthma (Fig. 4, S4 Table). These suggested the upregulation of *PTAFR* might serve as a biomarker for asthma severity.

**Figure 4.**
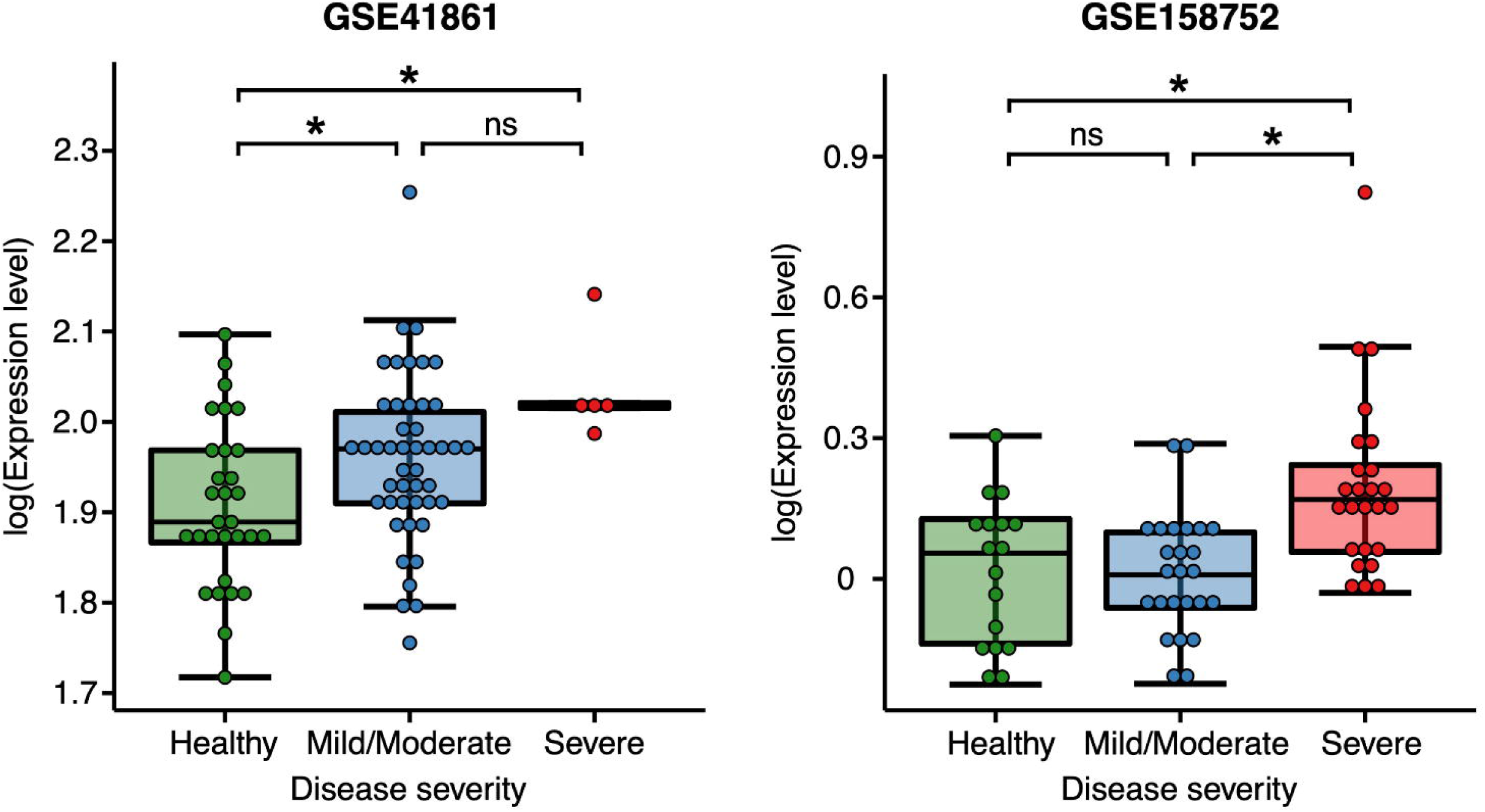
Upregulation of PTAFR was particularly observed in severe asthma. Expressions for PTAFR in the two asthma transcriptomics datasets having information on disease severity, i.e. GSE41861 and GSE158752, were displayed in box plots. *P < 0.05, ns: not significant. P < 0.05 is considered as statistically significant.

## Discussion

AI is revolutionizing the pharmaceutical industry by accelerating every single step of drug development, from the discovery of potential therapeutic targets(8), prediction of protein structures(43), generation of hit compounds(44), to the prediction of drug safety(45) and efficacy(46). The uses of AI in allergy prediction, diagnosis, and medicine were also documented(47, 48, 49). For instance, esophageal mRNA transcripts were analyzed by machine learning strategies (i.e., weighted factor analysis followed by random forest classification) to predict the probability of developing eosinophilic esophagitis during food allergy response with high sensitivity and specificity(50). Given the huge contribution of AI in the healthcare sector, this study applies the generative AI target discovery platform, PandaOmics, to reveal the commonly dysregulated biological processes and, more importantly, to identify targets with high therapeutic values in allergies.

PandaOmics has successfully demonstrated its power in retrieving high-quality therapeutic targets for multiple indications(9, 15, 51). We first evaluated the cellular processes that could serve as common disease-driving mechanisms across asthma, eczema, and food allergy. Immune dysregulation, the well-recognized underlying mechanism of allergies(52, 53), was revealed as one of the commonly dysregulated processes. Specifically, signalings of TRAF, NOD, and interferon were activated. These biological processes were directly associated with allergy development(54, 55, 56, 57). Mitochondrial malfunctioning, indicated by the dysregulation in pathways related to mitochondrial translation, mitochondrial biogenesis, and mitochondrial protein import, was identified as another disease signature. As triggered by allergens, oxidative stress, and reactive oxygen species (ROS) production were induced in the mitochondria, disturbing cellular bioenergetics and subsequently driving systemic inflammation(58, 59, 60). For instance, upon the oral challenge of peanut extract, the sensitized mice experienced reduced fatty acid oxidation, lower respiration rate, and increased ROS production(61). Antigen-dependent mast cell stimulation in allergies was also a consequence of disrupted mitochondrial dynamics(62). These collectively imply the alignment between our results and the published evidence.

Some groups of dysregulated pathways remain to be explored in allergy. For example, signaling by NTRKs was generally activated across the indications. Upon the binding of neurotrophins, activated NTRK receptors promote neuron growth, differentiation, and survival via the subsequent activation of downstream signaling pathways, i.e., MAPK, PI3K, and PKC(63). In a transcriptome associated study for pediatric asthma, NTRK1 and NTRK2 were the top differentially expressed genes(64). NT4/NTRK2 signaling also served as a cause of chronic airway hyperreactivity due to early-life allergen exposure(65). It is suggested that activation of NTRKs stimulated eosinophilic activity(66, 67). In view of the strong linkage between eosinophilic response and allergies(68), dysregulation of the NTRK pathways is worth additional effort in investigating its linkage with the allergic response.

In view of the successful retrieval of therapeutic targets with approved drugs for allergies, we demonstrated the power of PandaOmics in target discovery for allergic disorders. Three targets (*IL5*, *PTAFR,* and *RNF19B)* with potential therapeutic benefits to allergies were proposed in the study. All three targets were top-ranked in PandaOmics, with relatively consistent expression dysregulation profiles and high druggability. Both IL5 and PTAFR have launched drugs in the markets. It is worthwhile to conduct additional research to explore the possibility of repurposing those drugs for the treatment of allergic conditions. In addition, we observed the correlation between *PTAFR* expression and the severity of asthma. In the two datasets where *PTAFR* was significantly upregulated, the dysregulation of PTAFR is particularly detected in the samples with severe asthma, suggesting PTAFR might serve as an indicator of asthma severity. RNF19B is a novel target for allergic disorders that calls for novel drug development. We recommend experimental validation of these targets for developing or repurposing potential drug candidates to treat common allergic diseases.

In conclusion, we applied PandaOmics, the AI-driven target discovery platform, to identify a number of druggable targets that are common to asthma, eczema, and food allergy, facilitating the possible drug development of efficient and cost-effective therapy for allergy patients. This study also demonstrated the potential power of AI to aid in drug discovery and the development of efficient and affordable therapies for disease treatment.

## ACKNOWLEDGMENTS

We would like to thank Dr. Alex Zhavoronkov and Dr. Feng Ren for reviewing the manuscript and Dr. Xi Long for figure design.

## DATA AVAILABILITY

The original findings presented in the study are included in the article/supporting material, further inquiries can be directed to the corresponding author.

## FINANCIAL DISCLOSURE STATEMENT

The author(s) received no specific funding for this work.

## COMPETING INTERESTS

BHML, AS, MC, and FP are affiliated with Insilico Medicine, an artificial intelligence powered drug discovery & development company.

## Supplemental Information

**S1 Table.** Dysregulated pathways in asthma, eczema, and food allergy groups.

**S2 Table.** Expression profiles of *IL5*, *PTAFR*, and *RNF19B* in allergies-related bulk transcriptomics comparisons for target identification.

**S3 Table.** Dysregulated pathways associated with *IL5*, *PTAFR*, and *RNF19B* in allergies-related comparisons.

**S4 Table.** Association between the expressions of *IL5*, *PTAFR*, and *RNF19B* and disease severity in asthma-related bulk transcriptomics comparisons.

## Notes

### Summary of Updates

Affiliation of Andre Rayner was updated.

